# The impact of short-term forearm immobilization and acipimox administration on muscle amino acid metabolism and insulin sensitivity in healthy, young volunteers

**DOI:** 10.1101/2023.10.10.561668

**Authors:** Marlou L. Dirks, Tom S.O. Jameson, Rob C. Andrews, Mandy V. Dunlop, Doaa R. Abdelrahman, Andrew J. Murton, Benjamin T. Wall, Francis B. Stephens

## Abstract

The mechanisms underpinning short-term muscle disuse atrophy remain to be elucidated, but perturbations in lipid metabolism may be involved. Specifically, positive muscle non-esterified fatty acid (NEFA) balance has been implicated in the development of disuse-induced insulin and anabolic resistance. Our aim was to determine the impact of acipimox administration (i.e. pharmacologically lowering circulating NEFA availability) on muscle amino acid metabolism and insulin sensitivity during short-term disuse. Eighteen healthy individuals (age 22±1 years, BMI 24.0±0.6 kg·m^-2^) underwent 2 days of forearm cast immobilization with placebo (PLA; *n*=9, 5M/4F) or acipimox (ACI; 250 mg Olbetam; *n*=9, 4M/5F) ingestion four times daily. Before and after immobilization, whole-body glucose disposal rate (GDR), forearm glucose uptake (FGU, i.e. muscle insulin sensitivity), and amino acid kinetics were measured under fasting and hyperinsulinaemic-hyperaminoacidaemic-euglycaemic clamp conditions using arteriovenous forearm balance and intravenous L-[*ring*-^2^H_5_]phenylalanine infusions. Immobilization did not affect GDR but decreased insulin-stimulated FGU in both groups, but to a greater degree in ACI (from 53±8 to 12±5 µmol·min^-1^) than in PLA (from 52±8 to 38±13 µmol·min^-1^; *P*<0.05). In ACI only, fasting arterialised NEFA concentrations were elevated to 1.3±0.1 mmol·L^-1^ post-immobilization (*P*<0.05), and fasting forearm NEFA balance increased ∼4-fold (*P*=0.10). Forearm phenylalanine net balance tended to decrease following immobilization (*P*<0.10), driven by increases in phenylalanine rates of appearance (from 32±5 (fasting) and 21±4 (clamp) pre-immobilization to 53±8 and 31±4 post-immobilization; *P*<0.05) while rates of disappearance were unaffected and no effects of acipimox observed. Altogether, we show disuse-induced insulin resistance is accompanied by early signs of negative net muscle amino acid balance, which is driven by accelerated muscle amino acid efflux. Acutely elevated NEFA availability worsened muscle insulin resistance without affecting muscle amino acid kinetics, suggesting that disuse-associated increased muscle NEFA uptake may contribute to inactivity-induced insulin resistance but does not represent an early mechanism causing anabolic resistance.

## Introduction

Short periods of muscle disuse, e.g. during illness or recovery from injury, lead to rapid and substantial muscle atrophy, which is associated with negative consequences including a loss of muscle strength and function [1–4]. This loss of muscle mass is caused by negative net muscle protein balance, likely largely driven by impaired muscle protein synthesis in the fasting and postprandial states, the latter termed anabolic resistance [5]. We have recently shown that postprandial muscle amino acid uptake is reduced following 7 days of immobilization [6], suggesting that anabolic resistance might (partially) be caused by limited intramuscular amino acid availability following protein ingestion. In parallel with changes in muscle amino acid metabolism, disuse also leads to the development of muscle insulin resistance, i.e. a 30-40% reduction in insulin-stimulated skeletal muscle glucose uptake [1, 7–9], which we have previously demonstrated to be maximally developed within 2 days of removing muscle contraction [10, 11]. Disuse-induced muscle anabolic and insulin resistance are clearly due to a lack of contractile stimuli which otherwise maintain or increase muscle amino acid and glucose metabolism, but the underlying metabolic mechanisms are yet to be elucidated.

Perturbations in muscle lipid handling have been suggested to underpin the development of anabolic and insulin resistance during muscle disuse. We have previously demonstrated that a shift towards positive non-esterified fatty acid (NEFA) balance occurs across the forearm in response to ingestion of a mixed meal after 2 and 7 days of immobilisation, which corresponded with insulin [10] and anabolic [5, 6, 12] resistance. Presumably this positive balance results in lipid accumulation within the muscle during disuse. Indeed, changes in intramuscular diacylglycerol metabolism occur in the first 7 days of disuse [1, 13], and more prolonged disuse (>7 days) is associated with intramyocellular lipid (IMCL) accumulation [14], implicating altered muscle lipid handling as a locus of control for insulin and anabolic resistance during disuse. In support, we have previously shown that increasing plasma NEFA concentrations by experimental intravenous lipid infusion directly induces both insulin and anabolic resistance [15], and that a high-fat hypercaloric diet during 7 days of immobilization exacerbates the disuse-induced blunting of postprandial forearm amino acid balance [6]. Thus, this raises the question of whether preventing the immobilisation induced increase in forearm NEFA balance can reduce anabolic and insulin resistance and, ultimately, (partially) ameliorate the muscle deterioration associated with disuse.

To address this hypothesis, we performed a double-blind, randomized controlled study to investigate the impact of pharmacologically suppressing circulating NEFA availability and, therefore, muscle lipid accumulation during two days of forearm immobilisation on muscle amino acid metabolism and whole-body and muscle insulin sensitivity for the first time. We used four times daily administration of 250 mg acipimox, a nicotinic acid analogue that inhibits adipose tissue lipolysis for around 6 hours and improves insulin sensitivity [16–18], so that muscle NEFA uptake would be reduced throughout the entire immobilisation period. Measurements of muscle glucose, amino acid, and NEFA balance were performed using the arteriovenous-deep venous forearm balance technique [6, 10] in the fasting state and during a hyperinsulinaemic-hyperaminoacidaemic-euglycaemic clamp. This permitted us to directly measure insulin sensitivity in a controlled ‘postprandial’ steady-state, prior to and immediately after two days of forearm immobilization. In order to provide further insight into physiological mechanisms underlying any changes in anabolic sensitivity, intravenous L-[*ring*-^2^H_5_]-phenylalanine infusions were used in parallel to measure rates of forearm amino acid disappearance (Rd) and appearance (Ra).

## Methods

### Participants

Twenty-two healthy, young males and females were included in the present study. The participant characteristics of the final eighteen participants included in the study is depicted in **Table 1**. Prior to inclusion onto the study, participants attended the Clinical Research Facility (CRF) at the Royal Devon University Healthcare NHS Foundation Trust for a routine medical screening to ensure their eligibility to take part. Participants were excluded if they fulfilled one or more of the following criteria: age below 18 or over 40 y, BMI below 18.5 or over 30 kg·m^-2^, metabolic impairment (e.g. type 1 or 2 diabetes), hypertension, cardiovascular disease, chronic use of any prescribed over the counter pharmaceuticals or nutritional supplements, a personal or family history of thrombosis/epilepsy/seizures/schizophrenia, known allergies for any of the pharmacological treatments, any disorders in muscle or lipid metabolism, presence of an ulcer in the stomach or gut, severe kidney problems, and pregnancy. All participants were informed on the nature and risks of the experiment before oral and written informed consent was obtained. Height and weight were measured, and body composition was determined by Air Displacement Plethysmography (Bodpod; Life Measurement, Inc., Concord, CA, USA). The present study was approved by the NHS Wales REC4 Research Ethics Committee in accordance with the Declaration of Helsinki (version October 2013). This study was part of larger trial investigating the effects of pharmacological manipulations of substrate availability on muscle health during forearm immobilization, registered on clinicaltrials.gov as NCT03866512.

**Table 1:** Participants’ characteristics.

|  | **PLA (*n*=9)** | **ACI (*n*=9)** |
| --- | --- | --- |
| **Sex (M/F)** | 5 / 4 | 4 / 5 |
| **Age (y)** | 23 ± 2 | 20 ± 1 |
| **Height (cm)** | 175 ± 3 | 172 ± 2 |
| **Body mass (kg)** | 72.9 ± 3.1 | 68.6 ± 3.2 |
| **BMI (kg·m<sup>-2</sup>)** | 23.9 ± 0.7 | 22.6 ± 0.9 |
| **Body fat (% of body mass)** | 25.1 ± 3.1 | 21.9 ± 3.1 |
| **Lean mass (kg)** | 54.8 ± 3.8 | 53.5 ± 2.1 |
| **Systolic blood pressure (mm Hg)** | 114 ± 4 | 116 ± 4 |
| **Diastolic blood pressure (mm Hg)** | 65 ± 2 | 70 ± 3 |

All variables $P>0.05$ .

### Experimental overview

Following inclusion, participants visited the CRF for a baseline metabolic test day during which fasting and postprandial forearm glucose uptake (FGU) and amino acid kinetics were measured using the arterialized venous-deep venous (AV-V) forearm balance method. Participants attended the CRF for the application of a forearm cast, which signified the beginning of the 2-day immobilization period. During these 48 h, participants were randomized into receiving one of the following two pharmacological treatments in a double-blind manner: 250 mg acipimox (ACI) or placebo (PLA), all to be taken four times daily. During those same two days, participants were provided with a fully controlled eucaloric diet. Following two days of forearm immobilization, pharmacological treatment, and standardized nutrition, the metabolic test day was repeated. The forearm cast was removed following the final test day.

### Metabolic test day

At 08:00, after an overnight fast from 22:00, participants arrived at the CRF for the metabolic test day. For females not using hormonal/intrauterine contraceptives, both test days were scheduled on one of the first 10 days of their menstrual cycle, i.e. the follicular phase. For females on oral contraceptives both test days were conducted outside the stop week. Participants rested on the bed in a semi-supine position for the entire metabolic test day. Intravenous cannulas were placed 1) anterograde in an antecubital vein of the non-immobilized hand for intravenous infusions, 2) retrograde into a dorsal hand vein of the non-immobilized hand for arterialized venous blood sampling, and 3) retrograde into a deep-lying antecubital vein of the (to-be) immobilized arm to sample venous blood draining the forearm muscle bed [19, 20]. The cannulated hand (cannula 2) was placed in a heated (55°C) hand warmer. Following collection of a baseline venous blood sample, a primed (0.5 mg·kg body weight^-1^), continuous (0.5 mg·kg body weight^-1^·h^-1^) infusion of L-[*ring*-^2^H_5_]phenylalanine (CK Isotopes Ltd, Newtown Unthank, UK) was started for the duration of the test day (*t* = -150 min). Arterialised-venous (AV) and deep-venous (V) blood was sampled simultaneously five times between *t* = -30 and *t* = 0 min to measure fasting forearm muscle metabolism. Brachial artery blood flow of the (to-be) immobilized arm was determined by high-resolution ultrasound imaging in duplex mode (∼12 MHz, Apogee, 1000. SIUI, China) prior to every blood sample. Luminal diameter was imaged 5 cm proximal to the antecubital fossa for a 2 sec period. At the same anatomic location mean blood velocity was determined by integration of the pulsed-wave Doppler signal for a minimum of 8 cardiac cycles [21]. Semi-automatic analyses of captured files was done using Brachial Analyzer for Research, version 6.10.2 (Medical Imaging Applications LLC, Coralville, IA, USA, [22]).

At *t*=0 min, a hyperinsulinaemic-hyperaminoacidaemic-euglycaemic clamp was started to examine postprandial forearm muscle metabolism. Hyperinsulinaemic-euglycaemic clamps allow for repeated steady state forearm balance measurements allowing to detect a surplus effect of a potential intervention on top of the already large impact of immobilization (i.e. ∼40% decrease in both muscle glucose uptake and muscle protein synthesis following 7 days of limb immobilization [10, 23]), but are also regarded as the gold-standard technique to measure whole-body glucose disposal. This allows interpretation of effects on the local forearm level in the light of potential changes in whole-body glucose disposal, currently not possible when these techniques are used in isolation. Since hyperinsulinaemic-euglycaemic clamps lead to a suppression of circulating amino acids due to insulin-induced suppression of protein breakdown [31, 32], the use of intravenous amino acid co-infusion induces a steady state situation with postprandial amino acid concentrations. Therefore, the following intravenous infusions were started in the antecubital elbow vein: a primed (0-5 min: 128.2 mU·m^2^·min^-1^; 5-10 min: 71.8 mU·m^2^·min^-1^), continuous (from 10 min: 50 mU·m^2^·min^-1^) infusion of insulin (Actrapid, Novo Nordisk Ltd, Gatwick, UK) and a primed (0.46 mL·kg body weight^-1^) continuous (1.38 mL·kg body weight^-1^·h^-1^) infusion of 10% Primene (Baxter Healthcare Ltd, Northampton, UK) which was spiked with 7% L-[*ring*-^2^H_5_]phenylalanine to minimize plasma tracer dilution. A variable rate of 20% dextrose (Baxter) infusion was started in the same cannula. Every 5 min throughout the entire 3 h clamp a 0.5 mL blood sample was taken to determine blood glucose concentration, and the amount of glucose infused was altered to maintain euglycaemia at 5.0 mmol·L^-1^. Potassium chloride (0.3% KCl in 0.9% NaCl, Baxter) was infused in the (to-be) immobilized arm at a rate of 1 mL·kg body weight^-1^·h^-1^ to prevent insulin-induced hypokalaemia. The first twelve participants completed the study without issues. Thereafter unexplainable nausea and sickness occurred at the end of the clamp in two participants (of which one dropped out). The final four volunteers in the study received prophylactic metoclopramide hydrochloride (10 mg) intravenously at *t*=120 min to prevent these issues. Metoclopramide infusion did not affect any of the observed results. Every 30 min from the start of the clamp, brachial artery blood flow was measured and AV and V blood was sampled simultaneously (by two different investigators). During the last half hour of the clamp (i.e. between *t* = 150 and *t* = 180 min), five simultaneous AV and V blood samples were collected to measure insulin-stimulated forearm muscle metabolism. The same steady-state period was used to calculate the mean glucose disposal rate (GDR).

Forearm glucose uptake and forearm non-esterified fatty acid (NEFA) balance were calculated as the AV-V difference in glucose and NEFA concentrations, respectively, multiplied by brachial artery blood flow [24], as reported previously [10]. Forearm amino acid kinetics were calculated as described previously [6]. As forearm volume correlated well with body weight in our previous work ([6], Pearson’s correlation 0.779, *P*<0.001), and did not change with 7 days of forearm immobilization [6], in the present study we estimated forearm volume by multiplying body weight by 12.7 to use in the calculations for amino acid kinetics.

### Forearm immobilization

On the morning of the start of the 2-day forearm immobilization period, participants arrived at the CRF at 8:00 am to have a forearm cast fitted. Firstly, stockinette and undercast padding were applied to protect the skin. Next, a fiberglass (Benecast™, BeneCare Medical, Manchester, UK) cast was fitted to the forearm and hand to immobilise the wrist. This resulted in a cast which extended from 5 cm distal of the antecubital fossa to 2 cm proximal of the fingertips. Participants were provided with a sling and instructed to wear that during all waking hours to keep the hand elevated above the elbow. A waterproof cover was provided to keep the cast dry whilst showering. The immobilized arm was randomized and counterbalanced for arm dominance. Body weight was measured after application of the cast and this was repeated at the start of the second metabolic test day.

### Pharmacological treatment

During the two days of forearm immobilization, participants were randomly allocated to receive one of the following two pharmacological treatments in a double-blind manner: 250 mg acipimox (Olbetam, Pfizer Ltd, Sandwich, UK), or an inert placebo (containing microcrystalline cellulose, lactose, and magnesium stearate, manufactured by the Guy’s and St Thomas’ NHS Foundation Trust Pharmacy Manufacturing Unit). Treatments were prepared by the Royal Devon University Healthcare NHS Foundation Trust Clinical Trials Pharmacy and dispensed in opaque containers by a CRF research nurse blinded to treatment. Both treatments were orally ingested four times daily, i.e. at 8:00, 13:00, 18:00, and 23:00 (with the final dose on the second day taken at 22:00). Participants were instructed to take their treatment with water, and with/immediately after a meal or snack. Compliance was monitored via provided treatment logs, returned containers, and daily communication with study participants.

### Dietary intake

Prior to the immobilization period participants were instructed to keep a food diary for three consecutive days, including two weekdays and one weekend day. These food diaries were used to calculate habitual energy and macronutrient intake using the online licensed Nutritics software [25]. During the two days of forearm immobilization, participants received a fully-controlled eucaloric diet as described previously [10]. All meals and snacks were provided, whereas water and non-caloric drinks were allowed ad libitum. Energy requirements were individually calculated as basal metabolic rate (BMR via Henry equations [26]) multiplied by an activity factor (International Physical Activity Questionnaire, IPAQ; [27]). The diet was designed to provide 1.2 g protein·kg body weight^-1^·d^-1^, with a target macronutrient composition of 50-55 energy percent (en%) carbohydrate, 30-35 en% fat, 10-15 en% protein, and 2 en% dietary fibre. Compliance with the provided diet was assessed via completed 2-day food diaries, returned food containers, and daily communication with study participants.

### Sample analyses

Arterialized venous and deep-venous blood samples were collected for determination of whole-blood glucose, plasma amino acid concentrations and stable isotope enrichments, and serum insulin and NEFA concentrations. Therefore, one part of every sample (1 mL) was collected in a BD Vacutainer® fluoride/oxalate tube, rolled on a tube roller for 2 min to inhibit glycolysis, and subsequently analysed for whole blood glucose concentrations (YSI 2500 blood glucose analyser, Xylem Analytics UK, Tunbridge Wells, UK). A second part (5 mL) was collected in BD Vacutainer® SST II tubes, which were left to clot at room temperature for ≥30 min and then centrifuged at 2,500*g* at 4°C for 10 min to obtain serum samples. Arterialized serum samples were used to determine insulin concentrations (Human insulin ELISA kit, DX-EIA-2935; Oxford Biosystems Ltd, Milton Park, UK). Serum NEFA concentrations were measured spectrophotometrically in arterialized venous and deep-venous serum samples (FA115 kit, Randox Laboratories Ltd, Crumlin, UK). A third part of every sample (4 mL) was collected in BD Vacutainer® PST Lithium Heparin tubes and immediately centrifuged at 2,500*g* at 4°C for 10 min to obtain plasma samples. Plasma amino acid concentrations and L-[*ring*-^2^H_5_]phenylalanine enrichments were analysed using gas chromatography-mass spectrometry as described previously [6].

### Statistics

All data are expressed as means±SEM. Baseline characteristics between groups were tested using an independent samples *t*-test. Data were analysed using a Repeated Measures ANOVA with immobilization (pre vs post), prandial state (fasting vs clamp), and/or time point (during test day) as within-subjects factors, and treatment (ACI vs PLA) as between-subjects factor. In case of a significant interaction additional Repeated Measures ANOVAs were performed, with subsequent Bonferroni post hoc tests applied where necessary to locate individual differences. Statistical data analysis was performed using SPSS version 27.0 (IBM Corp, Armonk, NY, USA). Statistical significance was set at *P*<0.05.

## Results

### Participants and dietary intake

The two treatment groups did not differ in any baseline characteristics (**Table 1**) or habitual dietary intake (**Table 2**) prior to the start of the study. Three participants dropped out during the study: two because of cannulation issues on the first metabolic test day, and one because of issues with nausea and sickness. One participant in the acipimox group was excluded as both their whole-body glucose disposal and forearm glucose uptake at baseline were >2 SD greater than the rest of the population, despite being classified as recreationally active. The standardized diet consumed during forearm immobilization contained more energy than their habitual diet (*P*<0.05) due to absolute and relative increases in dietary carbohydrate and fibre content (both *P*<0.05), whereas alcohol intake was removed. Although relative protein content of the diet decreased when compared with habitual intake (*P*<0.05), absolute protein intake was unchanged due to an increase in energy intake. No differences were observed in dietary intake between groups (all *P*>0.05). During the two days of forearm immobilization body weight decreased from 73.8±3.0 to 73.4±3.1 kg in PLA and from 69.8±2.9 to 69.0±3.0 kg in ACI (*P*<0.05), with no differences between groups (*P*>0.05).

**Table 2:**
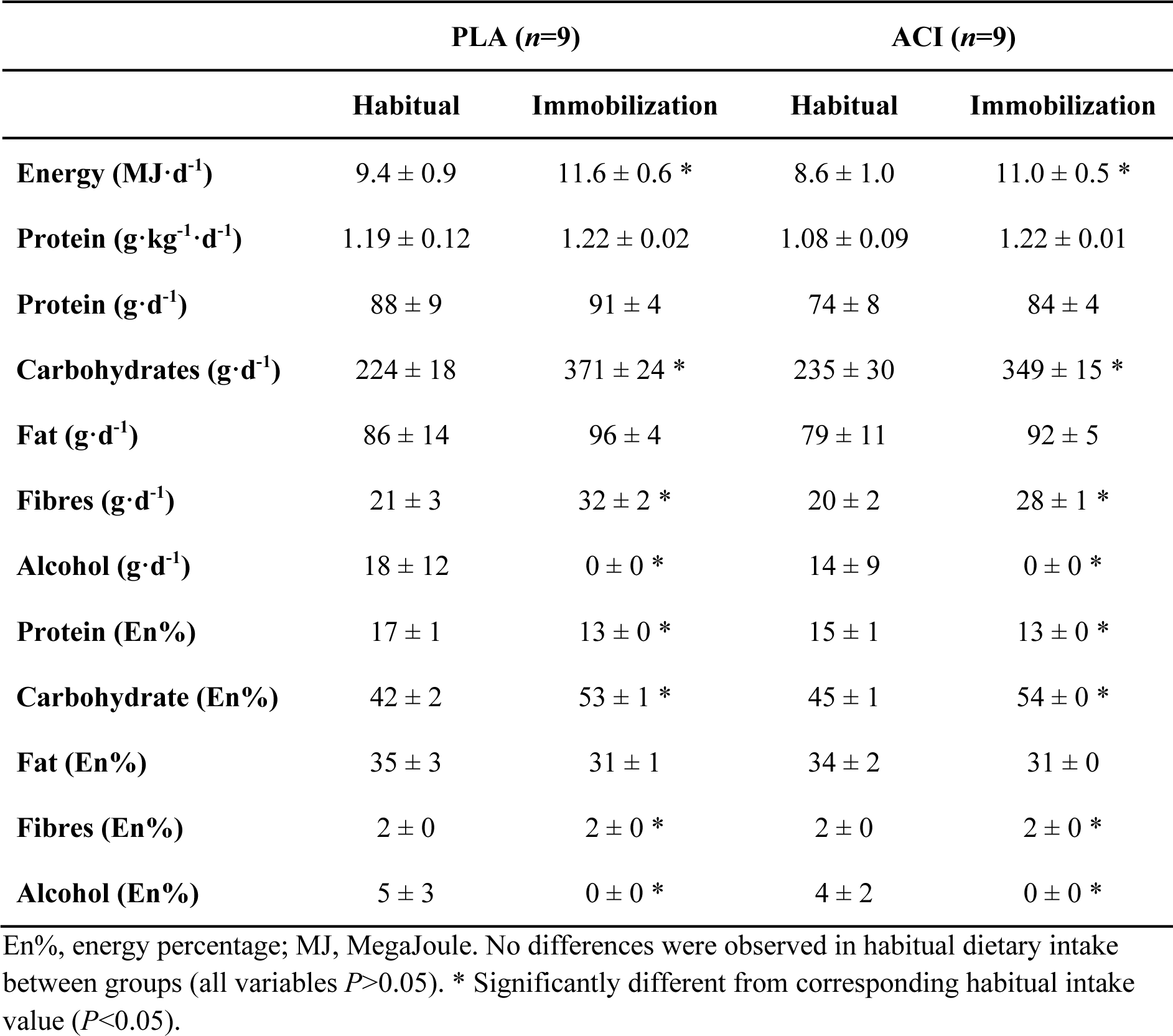
Dietary intake.

### Non-esterified fatty acids (NEFAs)

No differences in fasting serum NEFA concentrations were observed between groups prior to the study (*P*>0.05). Fasting arterialised serum NEFA concentrations increased with immobilization in both groups (*P*>0.05; **Figure 1A**+**B**), but to a greater extent in ACI (*P*<0.05). For arterialised serum NEFA concentrations during the clamp all main effects and interactions were statistically significant (all *P*<0.05). In both groups, hyperinsulinaemic-hyperaminoacidaemic-euglycaemia suppressed arterialised NEFA concentrations (*P*<0.05). The significant interaction effects were attributed to fasting NEFA concentrations being elevated following immobilization in ACI but not in PLA.

**Figure 1:**
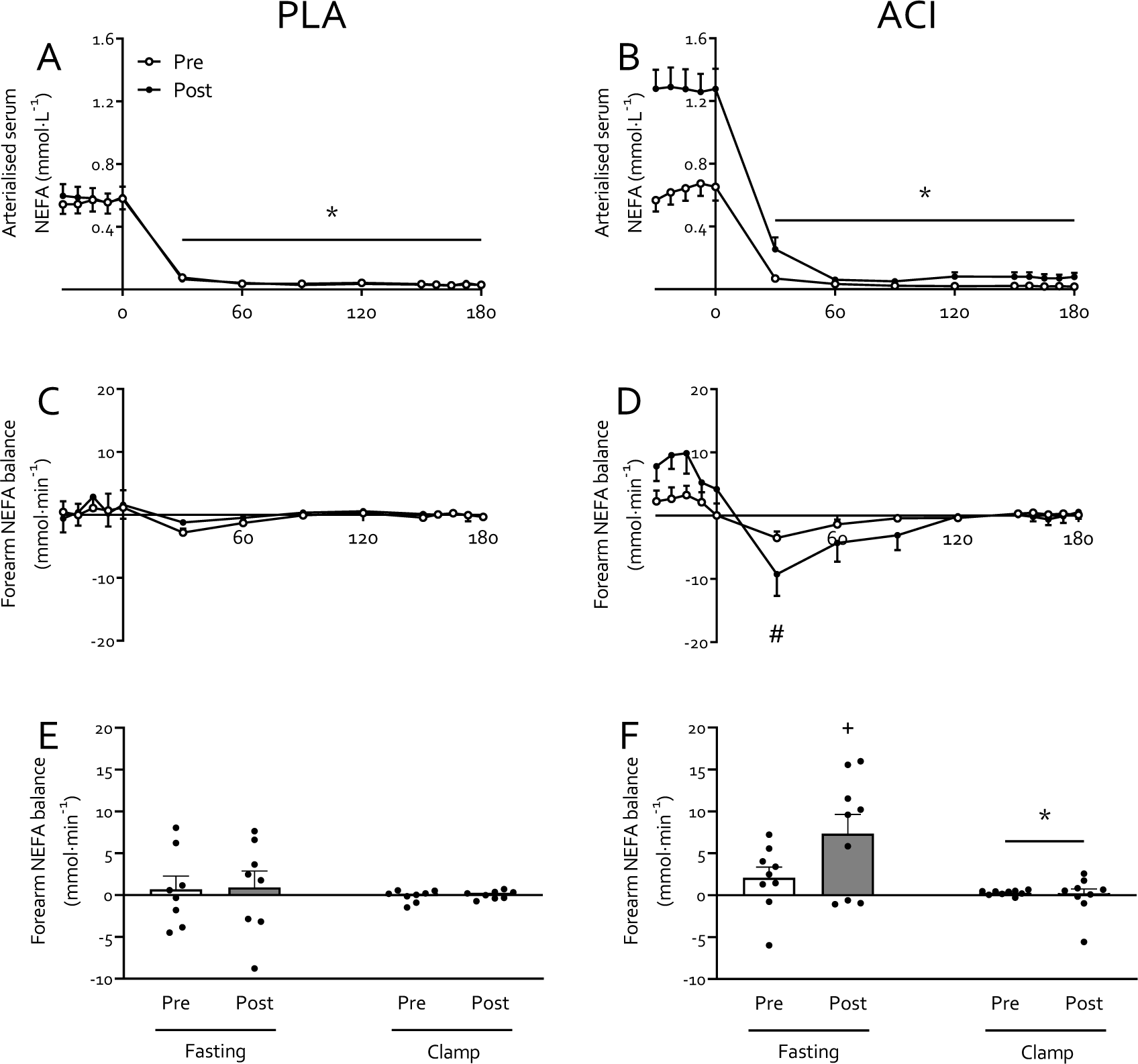
Non-esterified fatty acid (NEFA) concentrations and balance prior to and immediately following 2 days of forearm immobilization in healthy volunteers supplemented with placebo (*n*=8, left-hand panels) or acipimox (*n*=9, right-hand panels) four times daily. Panels **A** and **B** display arterialised NEFA concentrations in the fasting state and during the 3-hour hyperinsulinaemic-hyperaminoacidaemic-euglycaemic clamp. Forearm NEFA balance over time is displayed in **C** and **D**, with positive and negative values indicating a net uptake and release of NEFA in forearm tissues, respectively. **E** and **F** represent the average NEFA balance in the fasting state and during the clamp. * Significantly different from fasting (*P*<0.05). # Significantly different from t= -22.5 min (*P*<0.05). + Trend for difference from pre-immobilization value (*P*<0.10). Data are expressed as means±SEM.

Fasting forearm NEFA balance tended to increase with immobilization in ACI (*P*=0.10) but not in PLA (*P*=0.829; **Figure 1C**+**D**). Forearm NEFA balance demonstrated a time effect and time*treatment interaction (both *P*<0.05), which were attributed to a time effect and immobilization*time interaction (both *P*<0.05) in ACI only. Specifically, this was due to a lower forearm NEFA balance at 30 min following the start of the clamp when compared to the t = -22.5 min fasting value (**Figure 1D**). As a result, the average forearm NEFA balance was reduced during the clamp when compared to the fasting state (*P*<0.05), with no effect of immobilization (*P*>0.05) but a trend for overall higher values in ACI (*P*<0.10) and for a clamp*treatment interaction (*P*<0.10; **Figure 1F**).

### Whole-body insulin sensitivity

No differences were found on fasting blood glucose or serum insulin concentration, and glucose disposal rate (GDR), between PLA and ACI during the pre-immobilization test day (all *P*>0.05). Fasting blood glucose concentration decreased in both groups with immobilization (*P*<0.05) but to a greater extent in ACI (interaction: *P*<0.05), i.e. from 4.51±0.14 to 4.45±0.07 in PLA and from 4.42±0.12 to 3.95±0.10 mmol·L^-1^ in ACI. Fasting serum insulin concentration (**Figure 2A**) remained unchanged during forearm immobilization in both groups (*P*>0.05), with values being 11.0±0.8 and 9.4±1.0 mU·L^-1^ in PLA and 10.6±1.0 and 11.0±1.2 mU·L^-1^ in ACI on the pre- and post-immobilization test days, respectively. During both pre- and post-immobilization clamps, circulating serum insulin concentration peaked at 160±6 mU·L^-1^ at t = 60 min and averaged 145±5 mU·L^-1^ at the end of the clamps, with no differences between groups (*P*>0.05). GDR displayed no significant effects or interaction (**Figure 2B**; all *P*>0.05).

**Figure 2:**
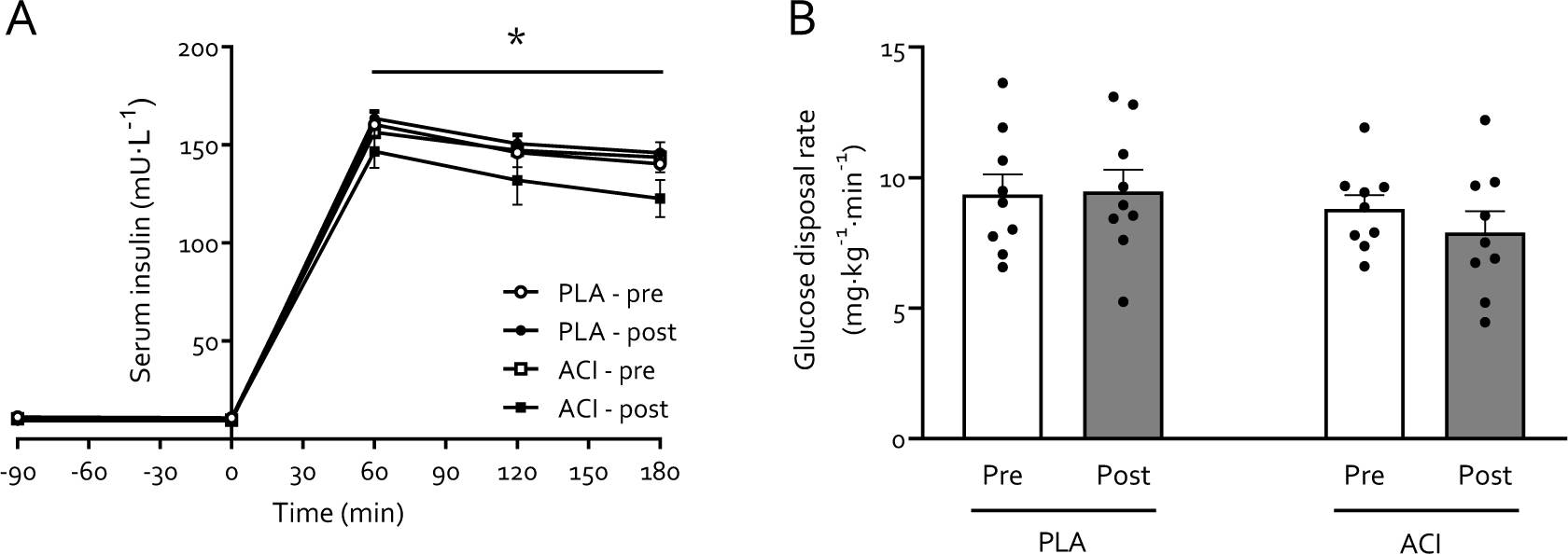
Serum insulin concentrations (**A**) during a 3-hour 50 mU·m^2^·min^-1^ hyperinsulinaemic-hyperaminoacidaemic-euglycaemic clamp in young healthy volunteers undergoing 2 days of forearm immobilization with placebo (PLA; *n*=9) or acipimox (ACI; *n*=9) supplementation. Panel **B** displays glucose disposal rates (GDR) during the final 30 min of the 3-hour clamp, representing steady-state conditions. Data was analysed using Repeated Measures ANOVAs. Data are expressed as means±SEM. * Significantly different from fasting concentrations (*P*<0.05).

### Muscle insulin sensitivity

Brachial artery blood flow (**Figure 3A-B**) increased with immobilization (*P*<0.05), but to a greater extent in ACI (interaction *P*<0.10) and particularly in the fasted state (*P*<0.05). Fasting forearm glucose uptake (FGU; **Figure 3C-F**) was not different between treatments on the pre-immobilization test day (*P*>0.05). FGU increased on average 3-fold from fasting during the hyperinsulinaemic-hyperaminoacidaemic-euglycaemic clamp on both pre-and post-immobilization test days (*P*<0.05). Two days forearm immobilization led to a reduction in both fasting and insulin-stimulated FGU (*P*<0.05). Based on a trend for an immobilization*treatment interaction (*P*=0.097) groups were analysed separately, and demonstrated an immobilization*clamp interaction (*P*<0.05) in ACI only. This implies participants in ACI were unable to increase FGU during the post-immobilization clamp when compared to the fasting state (*P*>0.05), whereas the insulin-stimulated state still led to increased FGU in PLA (*P*<0.05). In other words, ACI led to impaired insulin-stimulated FGU following 2 days of forearm immobilization, an effect confirmed by a significant difference between pre- and post-immobilization insulin-stimulated FGU (paired *t*-test, ACI: *P*<0.05, PLA: *P*>0.05). These findings occurred despite increased brachial artery blood flow on the post-immobilization test day in ACI only (*P*<0.05; **Figure 3B**).

**Figure 3:**
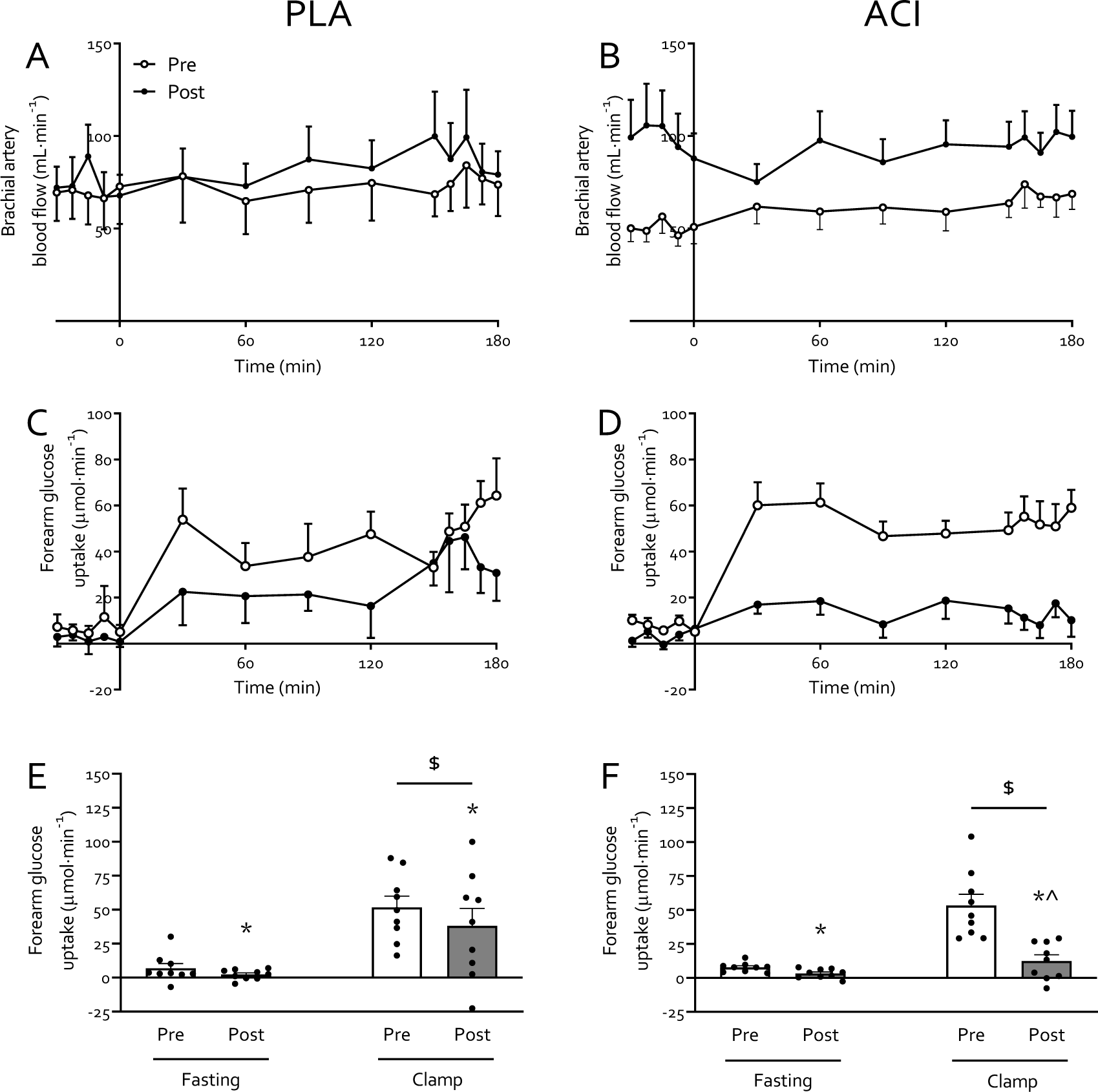
Muscle glucose uptake following 2 days of forearm immobilization with placebo (*n*=9; left-hand panels) or Acipimox (*n*=9; right-hand panels) supplementation in healthy young volunteers. Panels **A** and **B** display brachial artery blood flow, measured via Doppler ultrasound, which is used to calculate forearm glucose uptake (FGU; panels **C** and **D**) as a direct measure of muscle insulin sensitivity. Panels **E** and **F** represent the average FGU in the fasting state and during the clamp. * Significantly different from pre-immobilization. $ Significantly different from fasting (*P*<0.05). ^ Significantly different from pre-immobilization clamp value (*P*<0.05). Data are expressed as means±SEM.

### Amino acid concentrations and kinetics

Arterialised venous plasma leucine and phenylalanine concentrations increased from 125±4 and 47±1 to 272±7 and 98±3 µmol·L^-1^, respectively, during the transition from the fasting state to hyperinsulinaemic-hyperaminoacidaemic-euglycaemic clamp conditions (*P*<0.05), and were not affected by immobilization or treatment (both *P*>0.05; **Figure 4**). Plasma ^2^H_5_-phenylalanine enrichments (**Figure 4E**+**F**) increased moderately during the clamp (from 0.066±0.001 to 0.070±0.001 MPE in the fasting state and during clamp, respectively; *P*<0.05), but were not affected by immobilization or treatment (both *P*>0.05).

**Figure 4:**
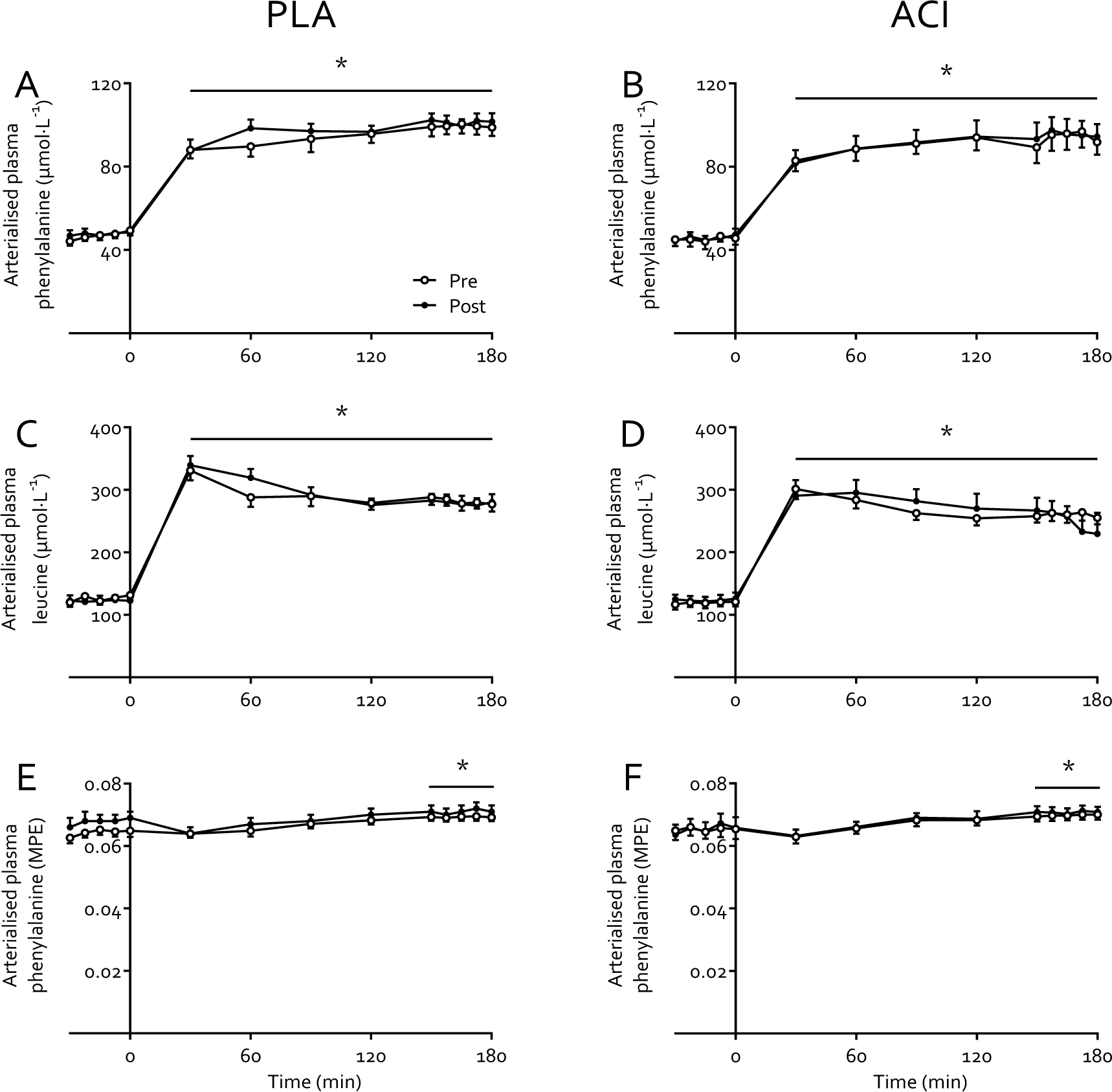
Arterialised plasma leucine concentrations (**A+B**), phenylalanine concentrations (**C**+**D**), and L-[*ring*-^2^H_5_]phenylalanine enrichments (**E**+**F**) before and immediately following 2 days of forearm immobilization in healthy young volunteers, in the fasting state (-30-0 min) and during a 3-hour hyperinsulinaemic-hyperaminoacidaemic-euglycaemic clamp (0-180 min). Participants were supplemented with placebo (*n*=9, left-hand panels) or acipimox (*n*=8, right-hand panels) whilst consuming a fully-controlled diet. * Significantly higher than fasting values (*P*<0.05). Data are expressed as means±SEM.

Forearm net balance (NB) of both phenylalanine (**Figure 5A**+**B**) and leucine **(5C+D**) switched from negative (-13±4 and -32±9 nmol·min^-1^·100 mL forearm volume^-1^, respectively) to positive (17±6 and 109±16 nmol·min^-1^·100 mL forearm volume^-1^, respectively; *P*<0.05) from fasting to clamp conditions. Immobilization decreased leucine forearm NB (*P*<0.05) and tended to decrease phenylalanine forearm NB (*P*<0.10), with no effect of treatment (*P*>0.05). Forearm phenylalanine rate of disappearance (Rd, **5E**+**F**) increased from 19±5 to 36±10 and from 40±6 to 54±10 nmol·min^-1^·100 mL forearm volume^-1^ in PLA and ACI during fasting and clamp conditions, respectively (*P*<0.05), but was not affected by immobilization. Forearm phenylalanine Rd was overall higher in ACI than in PLA (*P*=0.050), but no interactions were observed (all *P*>0.05). Forearm phenylalanine rate of appearance (Ra, **5G**+**H**) was suppressed during clamps (*P*<0.05) and was elevated following immobilization (*P*<0.05). Moreover, forearm phenylalanine Ra was overall higher in ACI than in PLA (*P*<0.05), with a tendency for a clamp*treatment interaction (*P*=0.080). Lastly, forearm leucine oxidation (**5I**+**J**) increased from -10±8 in the fasting state to 81±14 nmol·min^-1^·100 mL forearm volume^-1^ during the clamp (*P*<0.05) and was not affected by immobilization or treatment (*P*>0.05).

**Figure 5:**
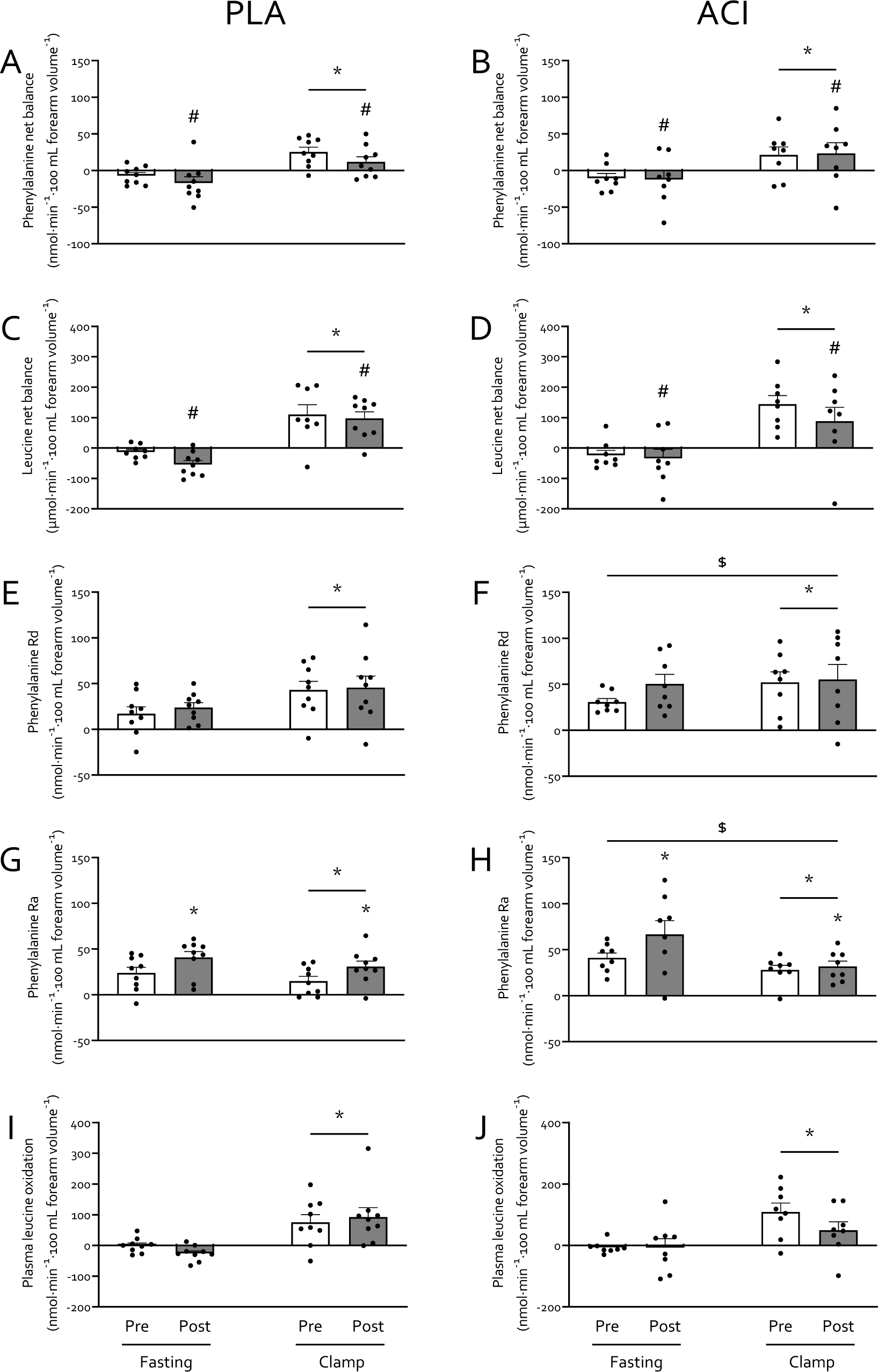
Amino acid kinetics prior to (white bars) and immediately after (grey bars) 2 days of forearm immobilization in healthy volunteers supplemented with placebo (*n*=9, left-hand panels) or acipimox (*n*=8, right-hand panels), in the fasting state and during the final 30 min of a 3-hour hyperinsulinaemic-hyperaminoacidaemic-euglycaemic clamp. Panels **A**+**B** and **C**+**D** represent leucine and phenylalanine net balance, respectively. panels **E**+**F** and **G**+**H** represent phenylalanine rate of disappearance (Rd; i.e. measure of muscle amino acid uptake) and rate of appearance (Ra; measure of muscle protein breakdown), respectively. Plasma leucine oxidation rates are depicted in panels **I**+**J**. * Significantly different from fasting (*P*<0.05). */# Effect of immobilization (* *P*<0.05; # *P*<0.10). $ Significantly higher than PLA group (*P*<0.05). Data are expressed as means±SEM.

## Discussion

The present study aimed to elucidate the role of positive muscle non-esterified fatty acid (NEFA) balance in immobilization-induced anabolic and insulin resistance by pharmacologically (via oral acipimox administration) supressing systemic NEFA availability. In contrast to our hypothesis, acipimox administration brought about a >2-fold elevation of fasting arterialised NEFA concentrations and >4-fold increase in fasting forearm NEFA balance during immobilization. As such, any effect(s) of repeated acipimox administration on circulating NEFA concentrations during 2 days of immobilization had either subsided, or were overridden by elevated NEFA availability, by the time forearm glucose and amino acid metabolism was determined. Nevertheless, this provided a unique scenario to investigate the role of increased NEFA availability on the anabolic and insulin resistance observed following disuse. Indeed, increased NEFA availability led to an exacerbated decrease in insulin-stimulated forearm glucose uptake in the absence of changes in whole-body glucose disposal. Moreover, we demonstrate for the first time that 2 days of forearm immobilization tends to decrease forearm phenylalanine net balance (NB) via an increased rate of phenylalanine appearance (Ra), suggesting increased amino acid efflux from muscle, while muscle amino acid uptake (Rd) was unaffected.

Periods of muscle disuse lead to the substantial development of insulin resistance, i.e. impaired insulin-stimulated glucose uptake, which occurs rapidly following the removal of muscle contraction [10]. Here we corroborate previous work [1, 7–9] by demonstrating that immobilization increases forearm NEFA balance ∼2-2.5 fold (Figure 1) and reduced forearm glucose uptake (i.e. direct measure of peripheral muscle insulin sensitivity) by ∼40% under hyperinsulinaemic-euglycaemic conditions, with hyperaminoacidaemic co-infusion (Figure 3). To understand the interaction between these disuse-induced perturbations of muscle lipid and glucose metabolism, we pharmacologically altered systemic lipid availability via oral acipimox administration. Acipimox is a nicotinic acid analogue that can acutely lower plasma NEFA concentrations by 60-75% for a 6 hour period [16, 28–31] via the inhibition of adipose tissue lipolysis, with repeated administration over several days previously being reported to improve insulin sensitivity and glucose tolerance in healthy normoglycemic [18, 32, 33] and insulin resistant [16–18] individuals. We assumed that four times daily administration of 250 mg acipimox during 2 days of forearm immobilization would lower plasma NEFA during the entire immobilisation period and, at least partially, prevent a positive muscle NEFA balance and subsequent muscle lipid accumulation during disuse. In contrast, however, we observed >2-fold higher serum NEFA concentrations following immobilization (Figure 1B), during measurements taken ∼10 h after the last acipimox dose. This is in line with a previously reported [29] ‘rebound’ effect of acipimox on lipolysis, suggesting any effect of lowering serum NEFA concentrations *during* the 2 days of immobilisation was obfuscated during the measurement period of forearm metabolism *following* immobilisation. Nevertheless, this provided a unique scenario to investigate the effect of an acute increase in NEFA balance on immobilisation-induced insulin resistance.

Elevated NEFA availability did not affect whole-body glucose disposal (Figure 2B), but participants receiving acipimox demonstrated a greater decrease in insulin-stimulated forearm glucose uptake during forearm immobilization than those supplemented with placebo (Figures 3C-F). We have previously demonstrated that a high-fat, hypercaloric diet (50% excess energy from fat) during 7 days of forearm immobilisation did not further exacerbate the positive NEFA balance or muscle insulin resistance induced by disuse [10]. Taken together with numerous reports demonstrating that acutely increasing circulating NEFA concentrations causes skeletal muscle insulin resistance, and that insulin resistance has plateaued by 24 hours of forearm immobilisation [11], this would suggest that acutely increasing circulating NEFA causes insulin resistance via a different, albeit transient, mechanism to a lack of contraction *per se* (e.g. Randle Cycle vs reduced GLUT4 translocation, respectively [13, 34]) during disuse. This does not rule out a role of muscle lipid accumulation in disuse-induced insulin resistance [35], but it has important implications for clinical scenarios where circulating lipids are elevated during physical inactivity and food intake requires adequate management, such as during critical illness [36].

To our knowledge, this is the first study to measure muscle amino acids metabolism following merely two days of limb immobilisation. In line with the rapid (i.e. within 2 days) development of insulin resistance with disuse, the present study also demonstrated a tendency for reduced muscle amino acid net balance (∼2-3 fold) under both fasting and clamp conditions during the same timeframe (Figure 5A), which is consistent with our previous observations following one week of disuse [6]. Although forearm muscle atrophy is not yet measurable via MRI so early into disuse [3], this negative amino acid balance is indicative of early muscle protein loss. Our experimental approach allowed us to estimate that immobilization reduced forearm net balance of all amino acids during the 30-min clamp steady state from 28.9 to 10.1 mg, representing net uptake of 0.5 and 0.2% of all amino acids infused, respectively. Interestingly, when using the assumptions that 12 h is spent in the fasted state daily, average forearm muscle mass is 0.6 kg [10, 20], and amino acids (as proteins) comprise 84% of human muscle tissue [37], this equates to a theoretical 0.73% daily muscle tissue loss. This is in line with what is observed in short-term leg immobilization studies in which muscle mass was quantified via MRI or CT [2, 38], but is approximately 2-3 fold greater than what is typically observed following short-term bed rest [4, 39]. This highlights the possibility of measuring early muscle protein loss, predicting subsequent measurable atrophy via imaging methods, directly in forearm muscles *in vivo,* which can act as an important early target in the development of effective interventional strategies.

Our measurements were conducted under tightly controlled hyperaminoacidaemic insulin clamp conditions, which resulted in elevated plasma amino acid concentrations comparable to peak plasma concentrations following ingestion of 35 g whey protein [40]. This approach obviated issues associated with applying a hyperinsulinaemic-euglycaemic clamp only to study ‘postprandial’ amino acid metabolism, whereby circulating amino acid concentrations decrease due to insulin-induced suppression of protein breakdown [41, 42]. By combining these clamp conditions with arteriovenous forearm balance measurements and an intravenous stable isotope tracer infusion we were able to demonstrate that the negative muscle protein balance observed after 2 days of immobilisation was not due to a reduced phenylalanine Rd, representing muscle amino acid uptake (Figure 5E). This contrasts our previous work in which we demonstrated a small, transient reduction in forearm phenylalanine Rd following 7 days of forearm immobilization in response to mixed meal ingestion [6]. This can potentially be explained by the clamp conditions being more anabolic than mixed meal ingestion, i.e. eliciting higher insulin and amino acid concentrations (Figures 2A and 4A, respectively). Although this requires confirmation in further research, this is in line with the potential for supraphysiological insulin concentrations to overcome age-related insulin resistance of protein metabolism [43].

Our data suggests that the removal of contraction *per se* rapidly induces insulin resistance but does not affect muscle amino acid uptake. This would fit with the different mechanisms and priorities of muscle contraction-mediated glucose and amino acid uptake (i.e. glucose is required as an immediate fuel source), and that the reduced amino acid uptake observed following 7 days of immobilization [6] is a physiological adaptation rather than a reduction in uptake capacity. Interestingly, as the Rd’s measured in the present work are similar to the peak Rd’s measured in response to mixed meal ingestion in our previous work (e.g. ∼50 nmol·min^-1^·100 mL forearm volume^-1^, [6]), this might indicate a maximal uptake capacity for amino acids in forearm muscle tissue. Nonetheless, the negative protein balance with 2 days of immobilisation appears to be due to an increase in phenylalanine Ra with immobilization (Figure 5G). Although it has been suggested that these amino acids may originate from increased muscle protein breakdown [23, 44] this has been debated [45, 46], and we recently showed that 2 days of leg immobilization did not affect fasting and postprandial muscle protein breakdown rates [12]. Instead, given amino acid oxidation was not affected by immobilization (Figure 5I; albeit in the face of lower energy demand), it is more likely that impaired muscle protein synthesis, which we have previously shown to occur over 2 days of limb immobilisation [12, 23], diverts excess amino acids to the circulation.

The increased NEFA balance and insulin resistance observed with immobilisation in the present study is in line with our previous work [6]. Specifically, we observed exacerbated immobilization-induced blunting of positive postprandial forearm amino acid balance when NEFA availability was further increased via 7 days of high-fat overfeeding [6]. Here we show that four times daily administration of 250 mg acipimox did not affect the immobilization-induced reduction in net balance of phenylalanine (Figure 5B) and leucine (Figure 5D), nor the phenylalanine Rd (Figure 5F) or Ra (Figure 5H), which is in contrast to the negative effect observed on glucose metabolism. Previous studies that have increased lipid availability via dietary means or intravenous infusion approaches have demonstrated reduced whole-body protein turnover and muscle amino acid efflux [47–49]. In agreement, we have previously demonstrated that acutely elevating NEFA availability combined with a hyperinsulinaemic-euglycaemic clamp almost completely supressed the muscle protein synthetic response to feeding [15]. This is difficult to reconcile with the present data, particularly given other studies have also demonstrated no effect or even increased muscle protein synthesis with elevated circulating NEFA [50, 51]. A possible explanation might be that providing energy from NEFA in the presence of amino acids and insulin creates a more favourable anabolic environment than amino acids and insulin alone, but that too much NEFA will lead to muscle lipid accumulation and subsequent impairments on anabolic signalling (e.g. suppressed 4E-BP1 phosphorylation, [15]). This might be particularly relevant in the presence of high insulin (Figure 2A), which will impair NEFA oxidation and release from muscle.

We conclude that rapid muscle insulin resistance observed with 2 days of forearm immobilisation is accompanied by early signs of reduced net muscle amino acid balance in both fasting and insulin-stimulated conditions, which is accompanied by an increase in amino acid efflux from muscle. Acutely elevating circulating NEFA availability with acipimox administration further decreased muscle glucose uptake but did not affect muscle amino acid metabolism. We therefore propose that increased muscle NEFA uptake with removal of muscle contraction may partly contribute to disuse-induced insulin-but not anabolic resistance. The latter thesis requires further investigation given acipimox administration in the present work may have reduced any detrimental effects of muscle lipid accumulation during immobilisation, thereby masking any effects on muscle protein metabolism by a subsequent ‘rebound’ effect of acutely elevated NEFA availability. The effect of lowering circulating NEFA on muscle deterioration during disuse remains to be investigated.

## Author contributions

MLD and FS designed the study. MLD, TSOJ, RCA, and MVD organised and carried out the clinical experiments. MLD, TSOJ, DRA, and AJM performed the laboratory analyses. MLD performed the statistical analyses. MLD, RCA, BTW, and FBS interpreted the primary data. MLD drafted, and RCA, BTW, and FBS edited and revised the manuscript. All authors approved the final version.

## Acknowledgements

We are grateful for the dedicated support from Claire Ball, Lauryn Domingos, Dr Mariana Coelho, Gráinne Whelehan, Freyja Haigh, Lindsay Wilkes, Silvia Balma, and Sarah Statton during data collection, and the Royal Devon University Healthcare NHS Foundation Trust Clinical Trials Pharmacy staff for their support with management and dispensing of study treatments.

